# Unveiling the influence of salt concentration on the different stages of the catalytic cycle of a halophilic enzyme

**DOI:** 10.1101/2025.01.14.632876

**Authors:** Gabriel Vallejos-Baccelliere, Sergio B. Kaufman

**Affiliations:** Laboratorio de Bioquímica y Biología Molecular, Departamento de Biología, Facultad de Ciencias, Universidad de Chile. Santiago, Chile; Departamento de Química Biológica, Facultad de Farmacia y Bioquímica. Universidad de Buenos Aires – IQUIFIB (UBA-CONICET). Buenos Aires, Argentina

**Keywords:** Halophilic enzyme, pre-steady-state kinetics, effect of salt in enzyme catalysis, enzyme-substrate charge repulsion

## Abstract

Enzymes from halophilic organisms have adapted to function in salt concentrations near saturation, making them an interesting model for studying the effect of salt on enzyme catalysis. The main insights into the effect of ionic strength on enzyme catalysis come primarily from enzymes with positively charged surfaces interacting with negatively charged substrates (e.g., ribonucleases), whose activity decreases at high salt concentrations. In this study, we investigated the effect of salt on the kinetics of Glucose-6-phosphate dehydrogenase (G6PDH) from the halophilic archaeon *Haloferax volcanii* (HvG6PDH), which has optimal activity conditions at concentrations exceeding 2 M KCl. The enzyme catalyzes the NAD+-dependent oxidation of G6P, a negatively charged substrate, and glucose, a non-charged substrate. Using steady-state kinetics, we determined that the enzyme follows an ordered-sequential kinetic mechanism, with NAD+ being the first substrate to bind and NADH being the last released product. Through steady-state kinetic experiments, we found that the main effect of salt is on the K_M_ for G6P, which decreased approximately 50-fold. For glucose dehydrogenase (glcDH) activity, the main effects were a 10-fold increase in k_cat_ and a roughly 10-fold increase in k_cat_/K_M_ for glucose. To analyze the effect of salt on the different stages of the catalytic cycle, we performed pre-steady-state experiments for both activities. We found that KCl did not affect the catalytic step in G6PDH activity, but it did increase the rate of catalysis in glcDH. Using a minimal model that accounts for substrate binding, chemical transformation, and product release, we determined that the main effect on G6PDH activity was an increase in the rate of G6P association. In contrast, for glcDH activity, the main effect was an increase in the rates of catalysis and product release. The results show that charge screening plays an essential role in the effect of salt on catalysis. Furthermore, it suggests differences in ion penetration to the active site between the two activities.

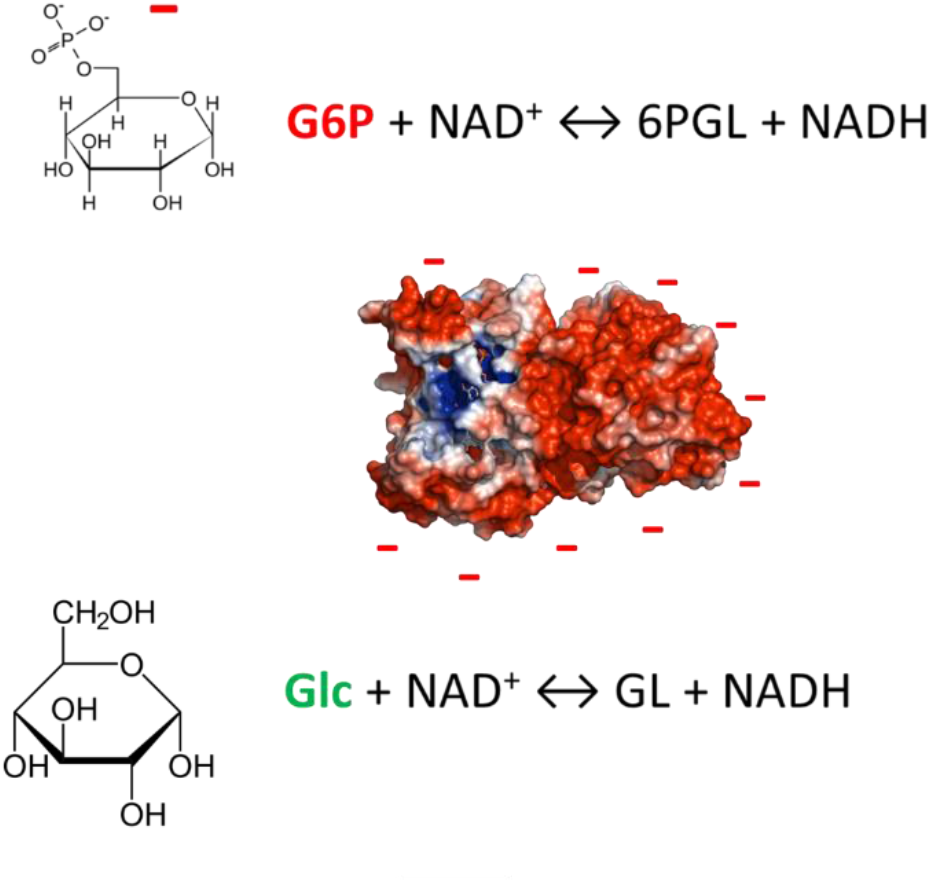

## Introduction

Halophilic organisms live in salt concentrations closer to saturation. During evolution, these organisms have developed different adaptative strategies, like the active expulsion of salt from the cytoplasm, the accumulation of organic osmolytes, or the accumulation of ions like potassium to avoid osmotic pressure stress. Consequently, proteins belonging to the latter group of organisms have developed physical-chemical properties to adapt to this functional context. These enzymes have been proposed as suitable candidates for biotechnological applications (Munawar et al. 2013, DasSarma & DasSarma 2015, Patel et al. 2015), and constitute an interesting model for studying the effect of salt on enzyme function.

The effect of salt concentration has been well described in functional macromolecular properties like solubility, stability (Jenkins 1998, Hyde et al. 2017, Song et al. 2007) and ligand binding, being the interaction with nucleic acids a subject with high theoretical development (Record et al. 1976, Overman et al. 1988, Cababie et al. 2019). However, the effect of salt on enzyme catalysis has not been studied to the same extent. In 2001, Park and Raines (Park & Raines 2001, 2003) developed a theoretical model to explain the decrease in enzyme activity caused by salt concentration in bovine pancreas ribonuclease. This protein has a positively charged surface and catalyzes the hydrolysis of a negatively charged linear substrate (RNA). By changing the enzyme surface charge in the vicinity of the active site and using RNAs with different lengths, they conclude that the main effect of salt would be a decrease in electrostatic attraction between the enzyme and the substrate (Park & Raines 2001, 2003). However, the validity of this theoretical model is restricted to enzymes whose activity decreases with salt concentration and whose surface charge is the opposite to the charge of the substrate (Park et al. 2003, Plantinga et al. 2008). It remains unclear whether this model can be applied to enzymes with other characteristics, like a different surface charge, or with non-linear polymeric substrates.

Halophilic enzymes have opposite characteristics. First, salt concentration produces an increase in their activity, reaching optimum values at molar concentrations (Mevarech et al. 2000). On the other hand, one common adaptive strategy in this kind of enzymes has been the development of a highly negatively charged surface (DasSarma & DasSarma 2015; Graziano & Merlino 2014; Jolley et al. 1997; Nath 2006). Studies have been reported for their stability, structure, and general kinetic properties (Jolley et al. 1997, Mevarech et al. 2000, Wright et al. 2002, Gonzalez-Ordenes et al. 2018), but a detailed kinetic study assessing the effect of salt in the different stages of the catalytic cycle has not been yet carried out.

A suitable model for studying the effects of salt concentration in enzyme activity is the glucose-6-phosphate dehydrogenase from the halophilic archaea *Haloferax volcanii* (HvG6PDH). This enzyme has no homology with the canonical G6PDHs from other organisms (Fuentes-Ugarte et al. 2021) and catalyzes the NAD^+^-dependent oxidation of G6P, producing 6-Phosphogluconolactone (6PGL) and NADH as products (Pickl et al. 2015). It can also catalyze the NAD^+^-dependent oxidation of glucose (glcDH activity) but with lower efficiency (Fuentes-Ugarte et al. 2021). The surface of this enzyme is enriched with acidic residues, while the active site cavity is enriched with basic residues, so the former has a net negative charge and the latter a positive charge in pHs above neutral (Fuentes-Ugarte et al. 2021). Since its substrate, G6P, has a negative charge, a repulsive interaction between the enzyme and this substrate is expected. Also, since glucose has no net charge, no repulsion is expected. On the other hand, NAD^+^ has both positive and negative moieties and, when transformed to NADH it loss a positive charge.

In this work, we performed a detailed analysis of the effect of salt concentration in the different stages of the catalytic cycle of HvG6PDH. First, we perform a comprehensive characterization of the effect of salt concentration on the enzyme using steady-state kinetic experiments to determine the kinetic mechanism of the enzyme and the effect of NaCl and KCl concentration on the kinetic parameters of both G6PDH and glcDH activities. To determine the effects of salt on the different stages of the catalytic cycle, we performed transient-state kinetic experiments for both activities at two KCl concentrations. We found that both activities are shifted toward product formation, the chemical transformation being the rate-limiting step in G6PDH and product release in glcDH. A joint analysis of stationary-state and transient-state kinetics results reveals that salt concentration exerts its effect in different stages of the catalytic cycle of each activity. We determine that the main effect of salt on G6PDH activity is an increase in the rate of association between G6P and the enzyme while in glcDH is an increment in catalysis, which is not affected in G6PDH activity. The results strongly suggest a role of charge screening and the entry of ions to the active site in the differential effect of salt in each activity.

## Results and discussion

### Kinetic mechanism determination

The kinetic mechanism of HvG6PDH was determined using steady-state kinetic experiments. Saturation curves for both substrates, G6P and NAD^+^, were obtained at different cosubstrate concentrations, NAD^+^ and G6P respectively (figure S1A-B), in 2 M NaCl. The Michaelis-Menten equation (equation 7) was fitted to each saturation curve and the apparent kinetic parameters k_cat_ and K_M_ for each substrate were estimated in each cosubstrate concentration. For both substrates, it was observed that k_cat_ behaves as a Michaelis-Menten Hyperbola and K_M_ remains constant as a function of cosubstrate concentration (Figure 1 A-D). This indicates that the kinetic mechanism occurs through the formation of a ternary E-NAD^+^-G6P complex. Otherwise, a Michaelis-Menten hyperbola would be expected for K_M_ vs. cosubstrate, which would have indicated a substituted enzyme mechanism.

**Figure 1.**
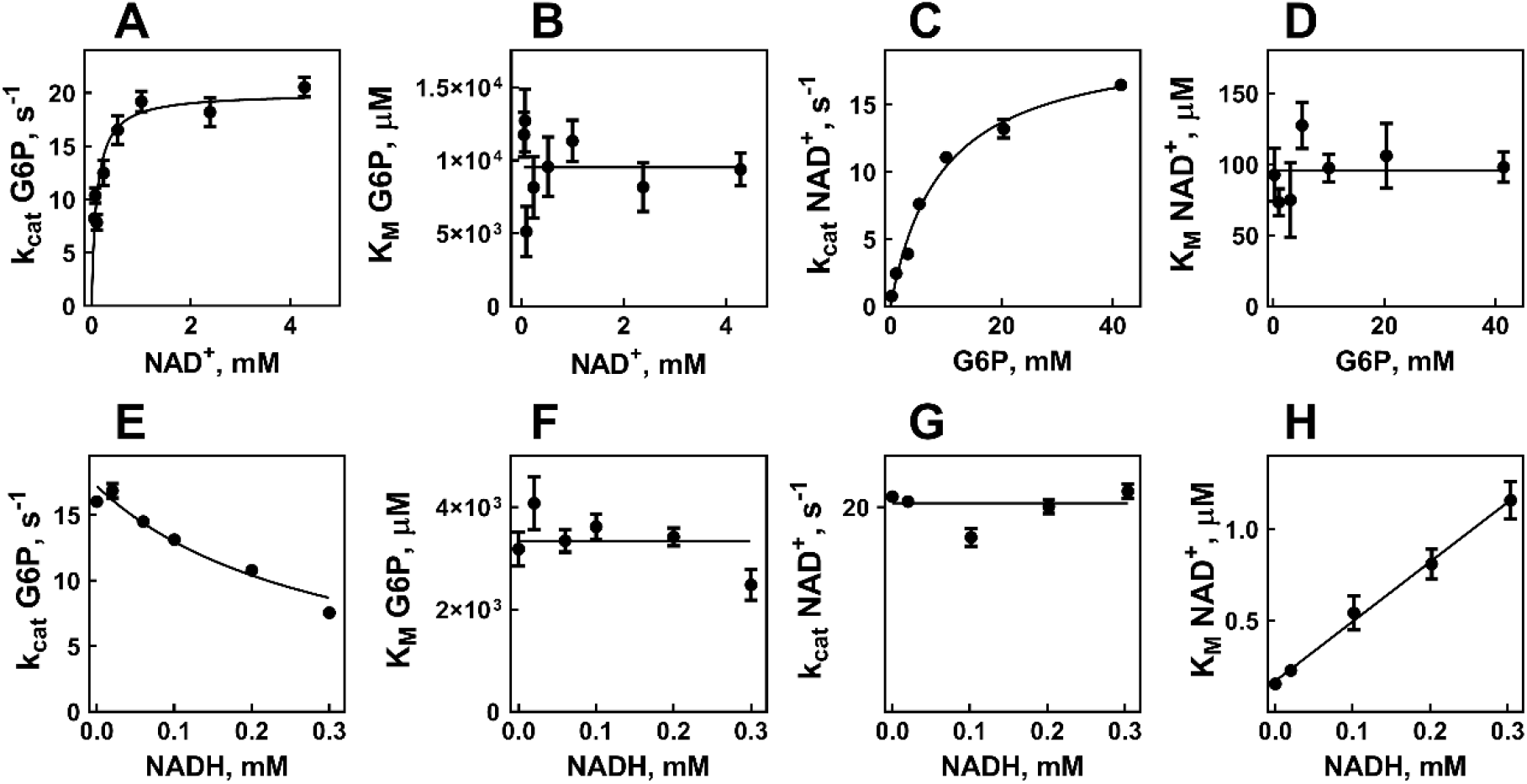
Steady-state kinetic experiments to determine the kinetic mechanism. k_cat_ for G6P at different NAD^+^ concentrations, a Michaelis-Menten hyperbola was fitted (continuous line). B) K_M_ for G6P at different NAD^+^ concentrations, the continuous line corresponds to the average value. C) k_cat_ for NAD at different G6P concentrations, a Michaelis-Menten hyperbola was fitted (continuous line). D) K_M_ for NAD^+^ at different G6P concentrations, the continuous line corresponds to the average value. E) k_cat_ for G6P at different NADH concentration, a hyperbolic decay with zero asymptote was fitted. F) K_M_ for G6P at different NADH concentrations, the continuous line corresponds to the average value. G) k_cat_ for NAD^+^ at different NADH concentration, the continuous line corresponds to the average value. H) K_M_ for NAD^+^ at different NADH concentrations, a straight line was fitted. Error bars correspond to the error in the estimation of parameter value by non-linear regression analysis.

To determine the order of substrate binding and product release, product inhibition studies were performed using NADH. Saturation curves were obtained for each substrate at different NADH concentrations, in 2 M KCl (Figure S1C-D). For G6P it was observed that k_cat_ vs NADH behaves as a hyperbolic decay with asymptote in zero (Figure 1E), while K_M_ remains constant (figure 1F). This is evidence that NADH is a non-competitive inhibitor regarding G6P. For NAD^+^, no effect was observed in k_cat_ (Figure 1G), while K_M_ vs NADH behaves as a straight line (Figure 1H). This is evidence that NADH behaves as a competitive inhibitor regarding NAD^+^. These results allow us to conclude that the kinetic mechanism of HvG6PDH corresponds to an ordered sequential, being NAD^+^ the first substrate to bind and NADH the last product released.

### Determination of the effect of salt concentration using steady-state kinetic experiments

To assess the effect of ions on the kinetics of HvG6PDH, first, we carried out steady-state kinetic experiments. For both G6PDH and glcDH activities, saturation curves for each substrate were performed in different KCl and NaCl concentrations (Figure S2). Since hyperbolic behaviors were obtained for each saturation curve, apparent kinetic parameters k_cat_ and K_M_ for each substrate were estimated as a function of salt concentration by fitting the Michaelis-Menten equation (equation 7). At this point, we use a phenomenological approach, so no theoretical model was presupposed to interpret the salt dependence of the kinetic parameters.

For G6PDH activity, the observed k_cat_ for NAD^+^ displayed a hyperbolic increase of 2.5 times when increasing NaCl concentration, and 6 times when increasing KCl (Figure 2A). On the other hand, a straight-line was found for 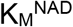 as a function of both NaCl and KCl concentrations (Figure 2B) with a slope slightly higher for KCl. Finally, a slight increase followed by an apparently hyperbolic decrease was observed 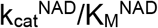 as a function of NaCl and KCl concentrations (Figure 2C). This behavior was consistent with a quotient of a hyperbola and a straight line.

**Figure 2.**
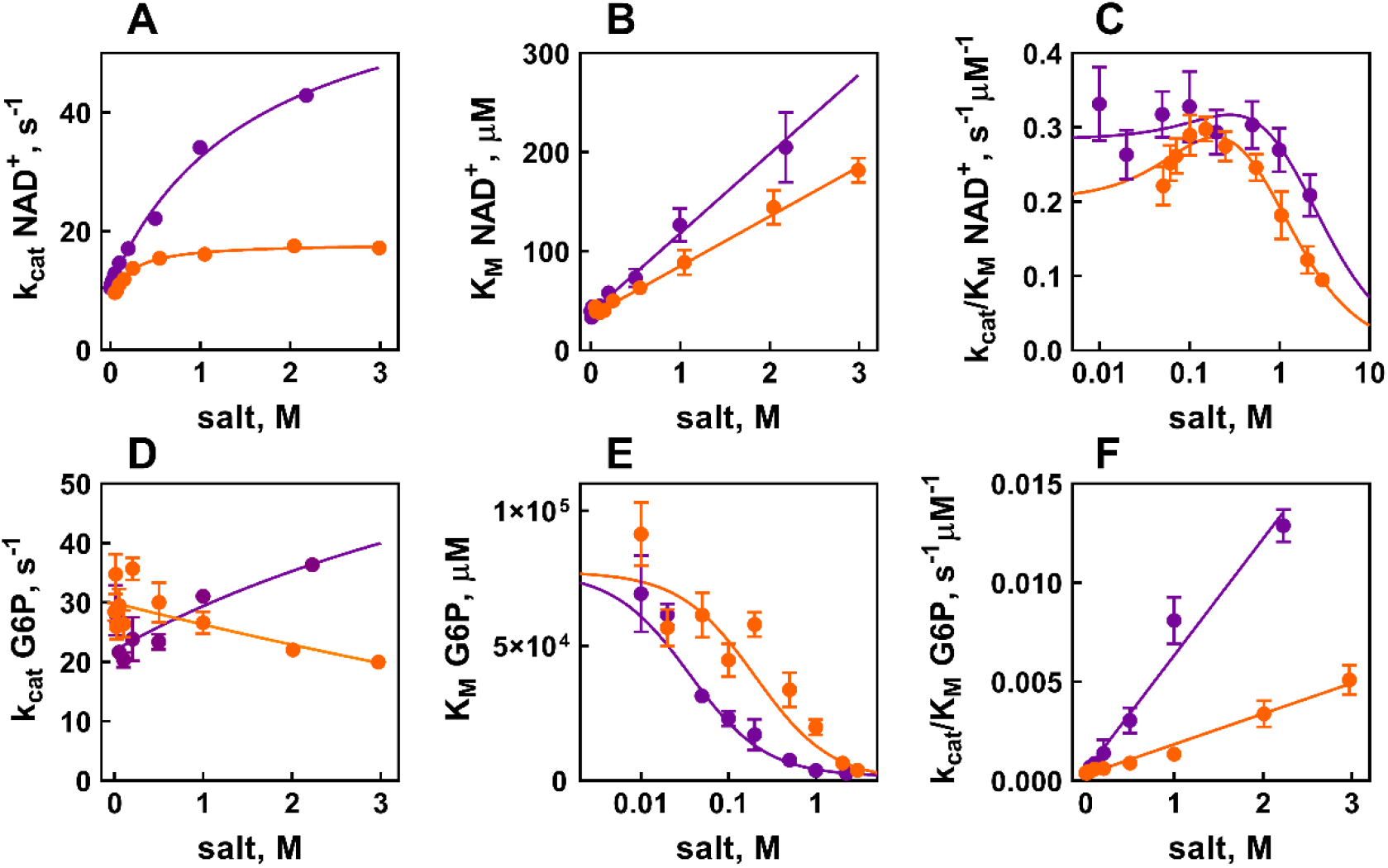
Analysis of salt concentration effect on G6PDH activity by steady-state kinetics. Secondary curves featuring the observed kinetic parameters k_cat_ (A), K_M_ (B) and k_cat_/K_M_ (C) for G6P and k_cat_ (D), K_M_ (E) and k_cat_/K_M_ (F) for NAD^+^ at different KCI 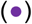 and NaCl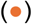 concentrations. The parameters were estimated by fitting the Michaelis-Menten equation to G6P and NAD^+^ saturation curves obtained at different fixed salt concentrations (figure S2). Error bars correspond to the error in the estimation of parameter value by non-linear regression analysis. Solid lines correspond to the fitted phenomenological equations; hyperbola intersecting the Y axis for (A), (D) and (E), straight line for B and F, and a quotient between a hyperbola and a straight line for (C).

For G6P, a slight 1.5 times hyperbolic decrease was observed for 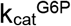 vs NaCl, while a hyperbolic increase of 2 times was observed as a function of KCl (Figure 2D). For 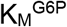, a pronounced hyperbolic decrease of more than 1 order of magnitude was observed as a function of both NaCl and KCl concentrations. The salt concentration needed to reach half of the saturation of the effect was bigger in NaCl than in KCl. For 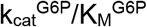 an approximately linear increase was observed as a function of both salt concentrations (Figure 2F), with an increase of approximately 40 times in the range of assessed concentrations of KCl and 10 times for NaCl.

HvG6PDH is also capable of catalyzing the NAD^+^-dependent oxidation of glucose. We analyzed the effect of NaCl and KCl concentration on glcDH activity by performing saturation curves for both NAD^+^ (Figure S3A-B) and glucose (Figure S3C-D) at different NaCl and KCl concentrations. A Michaelis-Menten Hyperbola was obtained for NAD^+^ saturation curves, and 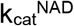 and 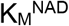 were estimated at different salt concentrations. A linear increase was observed for both 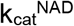 (Figure 3A) and 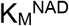 (Figure 3B) as a function of both NaCl and KCl concentrations. For 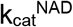 the effect of KCl was greater than the effect of NaCl, producing a 10-fold vs a 4-fold increment respectively in the range of assessed concentrations. This result contrasts with what was observed for G6PDH activity, in which this parameter has a hyperbolic dependence on salt with a lower increment. For 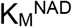, the effect of both salts was similar, producing an increase of approximately 3 times in both cases, not so different from what was obtained for the equivalent parameter in G6PDH activity. Finally, a saturation-like (hyperbolic) increase of 3 times was obtained for 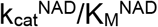 between 0 and 2 M salt (Figure 3C).

**Figure 3.**
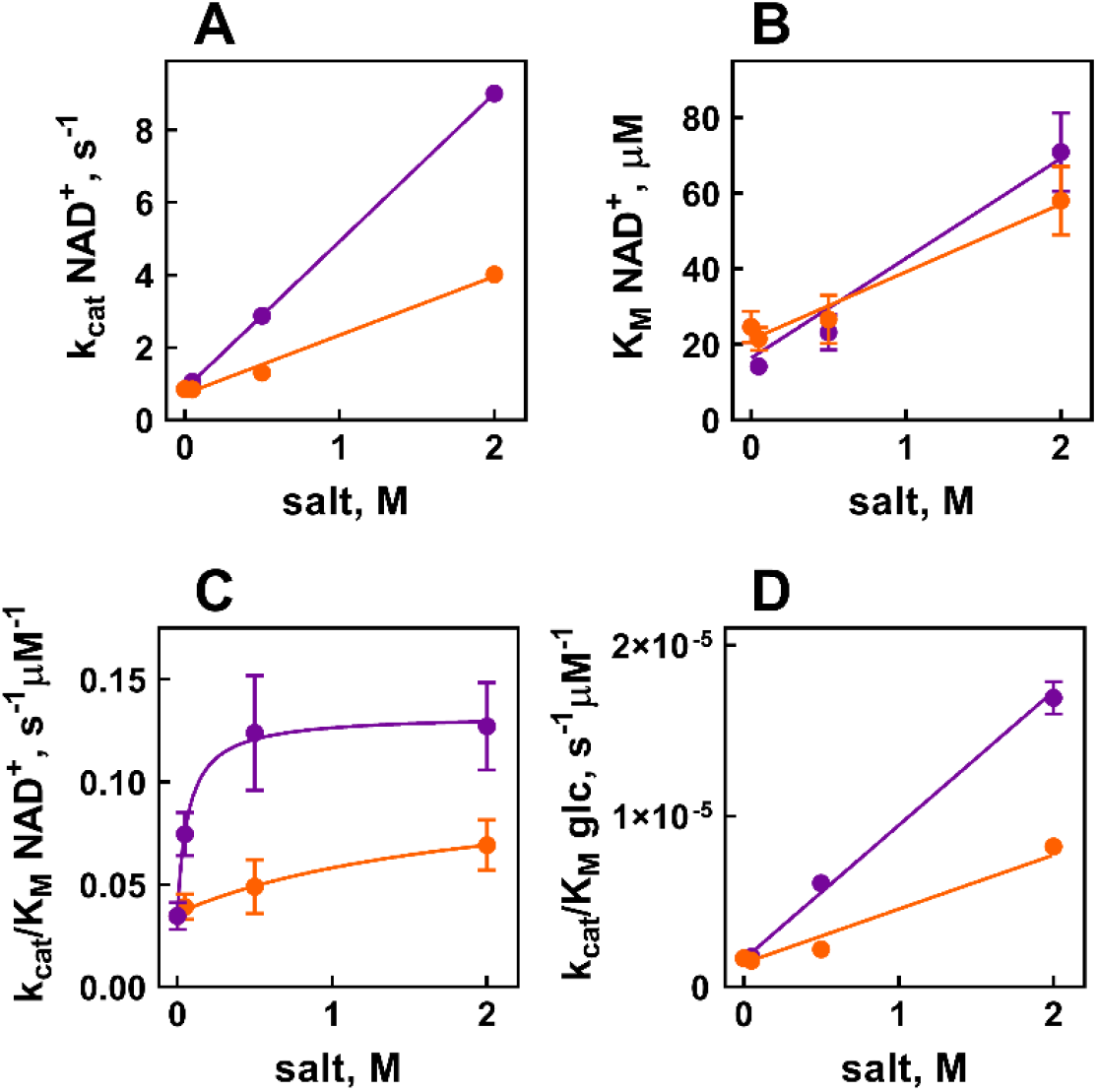
Analysis of salt concentration effect on G6PDH activity by steady-state kinetics. Secondary curves featuring the kinetic parameters k_cat_ (A), K_M_ (B) and k_cat_/K_M_ (C) for NAD^+^ and k_cat_/K_M_ (F) for glucose at different 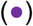 and 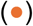 concentrations. The parameters were estimated by fitting the Michaelis-Menten equation to the NAD^+^ saturation curves and a straight line to the glucose curves (figure S3) Error bars correspond to the error in the estimation of parameter value by non-linear regression analysis. Solid lines correspond to the the best fitted phenomenological equations; a straight line for (A), (B) and (D), and a hyperbola intersecting the Y axis for (C).

Due to the low affinity of the enzyme for glucose, we didn’t reach saturation concentrations in the substrate curves (Figure S3C-D), so a straight line was fitted and the parameter 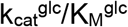 was estimated for each salt concentration (Figure 3D). A 5-fold linear increase in this parameter as a function of NaCl concentration was obtained, while KCl produced a linear increase of 10-fold. This increase was lower in comparison with the increase of 40-fold caused by salt concentration on 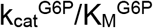 in G6PDH activity.

From the results, it is clear that salt concentration has contrary effects on both activities. In G6PDH the main effect is the dramatic decrease in 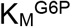, with lower effects in k_cat_ for both substrates. While in glcDH the main effect is the 10 times increase in k_cat_ NAD+. To address what are the bases of this differential effect of salt in HvG6PDH, it becomes necessary to assess the effect of salt in the different stages of the catalytic cycle of the enzyme.

### Transient-state kinetic experiments

To better understand the effect of salt concentration in both activities of HvG6PDH, and to have access to its effect on the different stages of the catalytic cycle, we perform time course experiments using a stopped-flow device and measuring NADH fluorescence (excitation at 340 nm and measuring emission intensity at 459 nm). A solution containing 2 µM NAD^+^ was mixed with solutions of equal volume containing increasing enzyme concentrations. Enzyme concentrations of 1 µM to 10 µM were assessed. Both activities were assayed under saturating sugar concentrations of 20 mM G6P and 0.2 M glucose respectively. The experiments were performed at 0.5 M and 2 M KCl.

For all time course curves for both activities and salt concentrations, no appreciable lag or burst phases were observed (Figure 4). Thus, a minimal model considering substrate binding, chemical transformation and product release steps was used to interpret the results.

**Figure 4.**
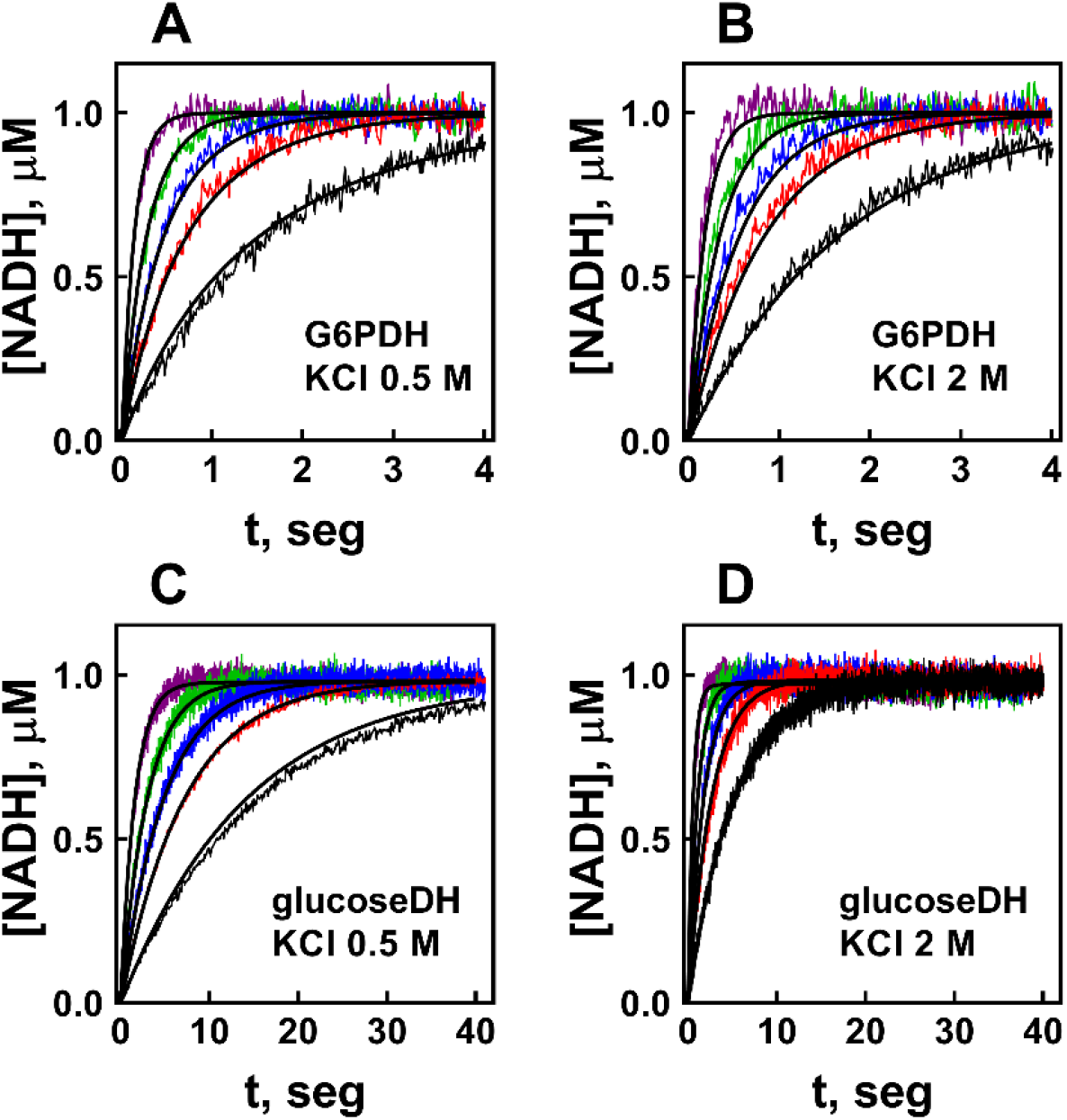
Time-course experiments. Time course experiments for G6PDH activity were performed at 0.5 M (A) and 2 M KCl (B) in 20 mM G6P, and for glcDH activity they were performed at 0.5 M (D) and 2 M KCl (E) in 0.2 M glucose. The experiments were carried out at different enzyme concentrations (**–** 1 µM, **–** 2 µM, **–** 3 µM, **–** 5 µM and **–** 10 µM). Solid lines represent simulations of the model indicated in scheme 1 using the parameters estimated by non-linear regression analysis shown in table 1.

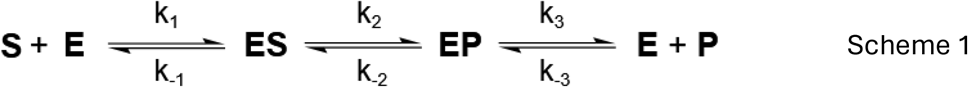

**Table 1.** Microscopic kinetic constants obtained by fitting the model represented in scheme 1. ‘e.e.’ denotes values for parameters obtained by stationary-state kinetic experiments

|  | G6PDH |  | glcDH |  |
| --- | --- | --- | --- | --- |
|  | 0,5 M KCl | 2 M KCl | 0,5 M KCl | 2 M KCl |
| $k_1, \mu\text{M}^{-1}\text{s}^{-1}$ | $0,9 \pm 0,1$ | $2,0 \pm 0,8$ | $0,111 \pm 0,007$ | $0,20476 \pm 0,00005$ |
| $k_{-1}, \text{s}^{-1}$ | $6 \pm 6$ | $125 \pm 63$ | $2,5 \pm 0,4$ | $1,25 \pm 0,09$ |
| $k_2, \text{s}^{-1}$ | $50 \pm 6$ | $55 \pm 6$ | $91 \pm 77$ | $365 \pm 3$ |
| $k_{-2}, \text{s}^{-1}$ | $< 0,007$ | $< 0,020$ | $84 \pm 77$ | $276 \pm 25$ |
| $k_3, \text{s}^{-1}$ | $62 \pm 73$ | -- | $4,4 \pm 0,7$ | $16,7 \pm 0,6$ |
| $k_{-3}, \mu\text{M}^{-1}\text{s}^{-1}$ | $68 \pm 79$ | $< 0,008$ | $0,0035 \pm 0,0004$ | $0,043 \pm 0,003$ |
| $K_S (k_{-1}/k_1), \mu\text{M}$ | $7,1 \pm 7,2$ | $63 \pm 41$ | $22,4 \pm 3,8$ | $6,1 \pm 0,4$ |
| $K_C (k_2/k_{-2})$ | -- | -- | $1,1 \pm 1,4$ | $1,3 \pm 0,1$ |
| $K_P (k_3/k_{-3}), \mu\text{M}$ | $0,9 \pm 1,5$ | -- | $1,250 \pm 245$ | $385 \pm 30$ |
| $k_{\text{cat}}, \text{s}^{-1}$ | **$28 \pm 37$** | **$48 \pm 240$** | **$2 \pm 2$** | **$9,3 \pm 0,5$** |
| $K_M, \mu\text{M}$ | **$34 \pm 43$** | **$80 \pm 350$** | **$31 \pm 28$** | **$48 \pm 3$** |
| $k_{\text{cat}}/K_M, \mu\text{M}^{-1}\text{s}^{-1}$ | **$0,8 \pm 1,5$** | **$0,6 \pm 4$** | **$0,07 \pm 0,1$** | **$0,19 \pm 0,01$** |
| $k_{\text{cat}} (\text{e.e.}), \text{s}^{-1}$ | **$24,4 \pm 0,2$** | **$42,5 \pm 0,1$** | **$2,87 \pm 0,07$** | **$9,0 \pm 0,2$** |
| $K_M (\text{e.e.}), \mu\text{M}$ | **$78 \pm 6$** | **$198 \pm 6$** | **$25 \pm 5$** | **$71 \pm 10$** |
| $k_{\text{cat}}/K_M (\text{e.e.}), \mu\text{M}^{-1}\text{s}^{-1}$ | **$0,31 \pm 0,07$** | **$0,22 \pm 0,03$** | **$0,12 \pm 0,02$** | **$0,13 \pm 0,02$** |

Because G6P and glucose were in excess with respect to NAD^+^ and the enzyme, it can be assumed that the binding of sugar after NAD^+^ is fast enough to justify this simplification. Because we didn’t reach enzyme saturation (i.e., single-turnover conditions), the analysis was performed without the use of pseudo-order or rapid equilibrium approximations.

We define K_S_ = k_-1_/k_1_ as the dissociation constant for the substrate, K_C_ = k_2_/k_-2_ as the equilibrium constant for the chemical step, and K_P_ = k_3_/k_-3_ as the dissociation constant for the product.

The stationary-state parameters deduced from this model are:

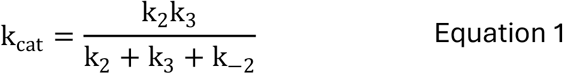

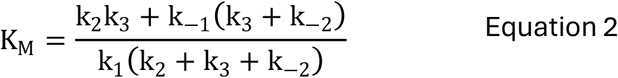

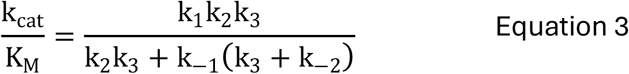

The space of possible values for the intrinsic kinetic constants was constrained by imposing an interval of allowed values to the k_cat_ and K_M_ deduced from the model (equations 1 and 2) around the empirical values for these parameters obtained from steady-state experiments. The model’s associated differential equations were numerically solved and fitted to the data using COPASI software. Simulations of the time-course curves are shown in figure 4.

For G6PDH activity, the chemical transformation is the limiting-rate step, and the equilibrium of the intermediary complexes is shifted towards product formation in both KCl concentrations, with k_2_ approximately 4 orders of magnitude over k_-2_ (table 1).

k_2_ is not affected by salt concentration; therefore, salt does not impact the catalytic step in this enzyme activity. However, KCl affects substrate affinity, producing a 9-fold increase in K_S_. This increase is due to an approximately 20-fold rise in k_-1_, while k_1_ only increases twice. This may be due to the salt enhancing repulsion between the substrate and the enzyme’s active site but decreasing it between the substrate and the enzyme surface, facilitating the encounter between both. Since k_2_ is unaffected by KCl, the observed increase in k_cat_ should be attributed to the salt effect on product release, i.e., an increment in k_3_ (table 1).

In glcDH activity, the rate-limiting step is product release, with k_3_ being about 20 times smaller than k_2_ in both KCl concentrations. In both cases, product association is negligible compared to its release, i.e., k_-3_ << k_3_. Contrary to G6PDH activity, in glcDH activity KCl increases NAD_+_ affinity, reducing K_S_ by approximately 4 times. This effect is due to a 2-fold increase in k_1_ and a halving of k_-1_. Interestingly, in this activity KCl does have an effect in the chemical step but impacts both k_2_ and k_-2_ in the same proportion, maintaining an equilibrium constant K_C_ around 1. The main effect of KCl on this activity is on product release, causing an increase of 4 times in k_3_ and decreasing product association approximately 10 times, resulting in a 3-fold decrease in K_P_. In this activity, the increase in 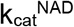 is due to increasing the rate of both first order processes occurring after NAD^+^ binding.

The charge screening of substrates seems to be one of the most important factors in the effect of salt in both enzymatic activities of HvG6PDH. This enzyme has a negative charged surface at the pHs used in the experiments (Fuentes-Ugarte et al., 2021). In the case of NAD^+^, it is expected that the negative charge of the pyrophosphate moiety primes over the positive charge of the nicotinamide. So, a decrease of repulsion between the enzyme and NAD^+^ by charge screening would be the explanation of the twice increment on the microscopic association constant for this substrate in both enzymatic activities. This is also consistent with the fact that NAD^+^ is the first substrate to bind. Furthermore, the active site cavity has a net positive charge (Fuentes-Ugarte et al., 2021), so charge screening would decrease electrostatic attraction between NAD^+^ and this site, increasing k_-1_. However, in glcDH activity, an opposite effect is observed, with a halving in k_-1_. This could be due to a difference in the entry of ions to the active site in each activity, probably because of the absence of the phosphate moiety in the sugar substrate.

Regarding the steps after substrate binding, in G6PDH activity there is no effect of salt, contrary to what happens in glcDH activity. This difference could also be due to differential penetration of salt to the active site. In glcDH activity, ions may be entering to the site of catalysis, but not in G6PDH activity. In this process, the presence or absence of phosphate moiety in the sugar substrate may also have a central role.

Charge screening can also explain the effect observed on product release stage. After the catalytic step, the dinucleotide substrate loss a positive charge, so it is expected that charge screening produces a decrease in the attraction between NADH and the positive-charged active site. This is consistent with the increment in product release rate (k_3_) observed in our results for glcDH activity. Additionally, a decrease in the repulsion between NADH and the enzyme surface by charge screening explains the increment in the rate of product association (k_-3_) in this activity. The effect cannot be evaluated in the case of G6PDH activity because it was not possible to model the product release step.

In the case of the effect of salt on sugar substrates, an indirect interpretation based on the combining analysis of transient-state and stationary-state kinetic results was performed. In the case of G6PDH activity, it can be assumed that salt concentration will have no effect on the catalytic step, and that the equilibrium is shifted toward product formation. Assuming that k_-2_ is negligible, 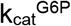 could be approximated to

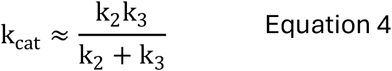

Assuming that k_2_ is not modified by KCl concentration, then the 2-fold increase on 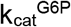 (Figure 2D) must be due necessarily to a 2-fold increase in the rate of product release (k_3_). Under the same assumptions,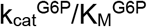 can be approximated to

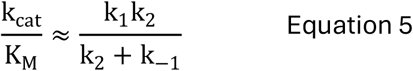

This parameter has a linear increase in the assessed interval of salt concentration (Figure 2F). Assuming that k_2_ is invariant, this effect can only be explained by a linear increase in k_1_ (it cannot be explained by a decrease in k_-1_, because to generate a linear increase in k_cat_/K_M_, it must necessarily take negative values after certain salt concentration). This linear increase in k_1_ can also explain the hyperbolic decrease observed for 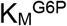.

These results allow us to conclude that the main effect of salt on G6PDH activity is an increment in the rate of G6P association, which can be explained by a decrease in the repulsion between this negatively charged substrate and the negatively charged enzyme by charge screening.

In the case of the effect on glucose the interpretation is more difficult because, due to the very high value of 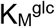, only 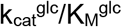 could be calculated as a function of salt concentration (Figure 3D). In this case, the only assumption that can be made following the pre-steady-state kinetics results is that k_2_ and k_-2_ are increased by salt in the same proportion, so K_C_ = k_2_/k_-2_ must be constat.

A very high value of K_M_ can be due to very low values in k_1_ or to very high values in k_-1_. Since glc has no net charge and hence no repulsion with the enzyme, it is likely that k_1_ is mostly associated with diffusion phenomena and the high value of 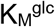 would be due to a very high value of k_-1_. Considering the data obtained in pre-steady-state kinetic experiments, it can also be assumed that k_-1_ >> k_2_.

Using K_C_ = k_2_/k_-2_, and k_-1_ >> k_2_, k_cat_/K_M_ can be expressed as

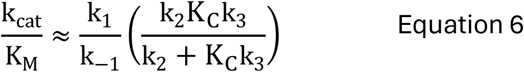

In the transient-state kinetic experiment results for NAD^+^, a which 4-times increment for k_2_ was observed, while for 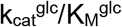, a 4-times increment was observed in the stationary-state kinetic experiments. So, the increment in k_2_ must be one of the main factors behind the increase in 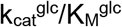.

The results conclude that the increase in enzymatic activity in G6PDH activity is mostly due to an increase in the rate of association of G6P, and, contrary, in glcDH is due mainly to an increase in catalysis and the stages occurring after substrate binding.

The effect of salt concentration on enzyme properties is a complex phenomenon. Each enzyme has unique features that can be affected by salt concentration. The cases previously studied regarding the effects of salt in catalysis were enzymes with positive surface charge and negatively charged linear substrates (RNAases) (Park & Raines 2001, 2003). In this work, we have studied an enzyme with a negatively charged surface, a positively charged active site cavity acting on substrates with different charge distributions. Other cases may exist in which the effect of salt could not be straightforwardly attributable. In the case we have analyzed, having a negatively charged surface corresponds to an adaptive property that halophilic proteins of this group of organisms have developed through evolution to function in high salt concentrations. But alternatives may exist. For example, Gonzalez-Ordenes et al. 2018 describe a non-canonical adaptation to high salt concentration in proteins from f halophilic archaea from the *Methanosarcinales* group. Due to the wide variety of possible cases, this work does not aim to develop a comprehensive theory on the effect of salt on enzyme catalysis. Using a detailed approach integrating steady-state and transient-state kinetics, we were able to differentiate the effect of salt in different stages of the catalytic cycle of the enzyme of both G6PDH and glcDH enzymatic activities. It remains to be seen how this effect may be occurring at the structural level. One probable explanation has to do with the fact that some interactions formed with the phosphate moiety of G6P will not be present when glucose is the substrate, leaving free space for the entrance of ions to the active site, so affecting the steps occurring after the enzyme-substrate complex formation.

## Materials and methods

### Expression and purification of HvG6PDH

Expression and purification of HvG6PDH was performed as described by (Fuentes-Ugarte et al. 2021). Briefly, the cDNA for HvG6PDH was cloned in the vector pET-TEV-28a with an N-terminal His-tag and the plasmid was inserted in E. coli BL21(DE3). Cells were cultured in L-B medium in 30 μg/ml Kanamicine at 37 °C until an OD^600^ of 0.8 -1. Protein expression was induced using 1 mM IPTG incubating overnight at 37 °C with continuous agitation. The cells were sedimented by centrifugation at 6,000 g for 20 minutes and resuspended in lysis buffer (2 M KCl, 50 mM Tris–HCl, pH 7.8, 1 mM EDTA, 5 mM 2-mercaptoethanol, and 0.1% v/v Triton x-114). Cell lysis was performed by sonication with 15 pulses of 20 s at 40 % amplitude separated with pauses of 1 minute.

The lysate was centrifuged at 38,000 g for 20 minutes, and the insoluble fraction was recovered and resuspended in solubilization buffer (8 M urea, 50 mM Tris–HCl, pH 7.8, 1 mM EDTA, and 50 mM dithiothreitol) in a volume corresponding to 2.5 % of the original culture volume and incubated for 1 h at 37 °C with permanent mild agitation. The solution was centrifuged at 38,000 g for 30 minutes and the supernatant was recovered. Proteins were refolded by a 20-folddilution in renaturation buffer (2 M KCl, 50 mM Tris–HCl, pH 7.8, and 5 mM 2-mercaptoethanol), incubating at 4°C overnight with permanent agitation. The refolded proteins were centrifuged at 38,000 g for 20 min, the supernatant was concentrated using a vivaflow filtration device (Sartorius) and loaded on a 5-ml HisTrap-HP column (GE Healthcare) previously equilibrated with renaturation buffer. The column was washed with 10 volumes of renaturation buffer supplemented with 50 mM Imidazole. Elution was performed with a 50 ml linear gradient from 50 to 250 mM imidazole. Fractions with G6PDH activity were pooled. Protein purity was evaluated by sodium dodecyl sulfate polyacrylamide gel electrophoresis (SDS-PAGE), for which the samples have to be previously precipitated with 20% trichloroacetic acid to remove the excess of KCl.

### Steady-state kinetics experiments

All experiments were performed in Tris-HCl pH 8.5 100 mM and different NAD^+^, G6P, NaCl and KCl concentrations. To control protein stability during the assay, selwyn tests and incubation experiments were performed. The reaction was started adding G6P or NAD^+^ after a pre-incubation of the enzyme for 5 minutes in the reaction conditions.

G6PDH and glcDH enzymatic activities were measured following NADH production by recording Abs^340^ considering an extinction coefficient of 6.22 mM^-1^cm^-1^. The experiments were performed in 96-well plates (Model 269 620; Nunc, Rochester, NY, USA) using a Synergy 2 spectrophotometer (BioTek, Winooski, VT, USA). Initial velocities were obtained by linear regression using Excel or GRAPHPAD PRISM software (Dotmatics, Boston, MA, USA). Initial velocity measurement was expressed in U/mg, where U corresponds to µmol/min.

Saturation curves for each substrate were obtained at different cosubstrate, NADH, KCl or NaCl concentrations and the Michaelis-Menten equation (equation 7) was fitted to the data of initial velocity vs substrate by non-linear regression using excel or GRAPHPAD PRISM software, and kinetic parameters k_cat_ and K_M_ were obtained as a function of cosubstrate or salt concentration. For k_cat_ calculations, a molecular mass of 30 KDa was considered for the protein (Fuentes-Ugarte et al. 2021).

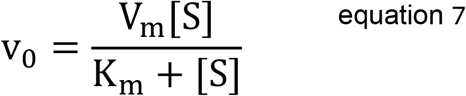

### Rapid kinetics experiments

All experiments were performed at 25 °C in Tris-HCl pH 8.5 100 mM, dithiothreitol 2 mM and 0.5 M or 2 M KCl. For G6PDH a constant concentration of 20 mM G6P was used, while for glcDH a concentration of 0.2 M glucose was used.

The experiments were performed using a rapid mixing stopped-flow accessory (RX2000 Applied Photophysics) coupled with a spectrofluorometer Jasco FP-8300. Reactions were initiated by mixing equal volume of a solution containing NAD^+^ 2 µM with a solution containing the enzyme at various concentrations from 2 µM to 20 µM. NADH fluorescence was measured by exciting at 340 nm and recording emission intensity at 459 nm. NADH concentrations were determined using a NADH calibration curve and correcting the measures by subtracting the fluorescence emission of both the protein and NAD^+^ in these conditions.

“A minimal model (Scheme 1) was used. The model was numerically solved and fitted to the time-course data using COPASI software. The search for values of the intrinsic kinetic constants (k_1_, k_-1_, k_2_, k_-2_, k_3_, k_-3_) was restricted by limiting the possible values of these constants to a defined interval of k_cat_ and K_M_ deduced from the model as a function of the intrinsic kinetic constants (equations 1 and 2). This interval was defined by values obtained from steady-state kinetic experiments for the parameters k_cat_ and K_M_

## Abbreviations

PNAD^+^: Nicotinamide adenine dinucleotide oxidized form.
NADH: Nicotinamide adenine dinucleotide reduced form.
G6P: glucose-6-phosphate.
G6PDH: G6P dehydrogenase.
glcDH: glucose dehydrogenase.
HvG6PDH: G6PDH enzyme from *Haloferax volcanii*.

## Author contributions

GVB led the research, designed and performed all the experiments, performed data analysis, developed kinetic models, and wrote the manuscript. SBK developed kinetic models and performed numerical solution and non-linear regression analysis.

## Acknowledgements

This work was supported by Fondo Nacional de Desarrollo Científico y Tecnológico (Fondecyt Grant 3210758) from the National Research and Development Agency (ANID), Chile. GVB would like to express acknowledgement to Dr. Victoria Guixé, Dr. Víctor Castro-Fernandez and Dr(c). Nicolás Fuentes-Ugarte from Laboratorio de Bioquímica y Biología Molecular for their support and contributions to this work. GVB also thanks Dr. Jorge Babul from Universidad de Chile for providing the necessary resources to perform the research.

## References

Cababie L, Incicco J, González-Lebrero R, Roman E, Gebhard L, Gamarnik A, Kaufman S (2019) Thermodynamic study of the effect of ions on the interaction between dengue virus NS3 helicase and single stranded RNA. Sci Rep. 22;9(1):10569.

DasSarma S & DasSarma P (2015) Halophiles and their enzymes: Negativity put to good use. Curr Opin Microbiol. 25: 120–126.

Fuentes-Ugarte, N., Herrera, S. M., Maturana, P., Castro-Fernandez, V., & Guixé, V. (2021). Structural and Kinetic Insights Into the Molecular Basis of Salt Tolerance of the Short-Chain Glucose-6-Phosphate Dehydrogenase From Haloferax volcanii. Frontiers in microbiology, 12, 730429.

Gonzalez-Ordenes F, Cea P, Fuentes-Ugarte F, Muñoz S, Zamora R, Leonardo D, Garratt R, Castro-Fernandez V, Guixé V. (2018) ADP-Dependent Kinases From the Archaeal Order Methanosarcinales Adapt to Salt by a Non-canonical Evolutionarily Conserved Strategy. Front Microbiol. 9:1305.

Graziano G, Merlino A (2014) Molecular bases of protein halotolerance. Biochim Biophys Acta. 1844(4):850–8.

Hyde A, Zultanski S, Waldman J, Zhong Y, Shevlin M, Peng F (2017) General Principles and Strategies for Salting-Out Informed by the Hofmeister Series. Org. Process Res. Dev. 21, 9, 1355–1370

Jenkins W (1998) Three solutions of the protein solubility problem. Protein Sci. 7(2): 376– 382.

Jolley K, Russell R, Hough D, Danson M (1997) Site-directed mutagenesis and halophilicity of dihydrolipoamide dehydrogenase from the halophilic archaeon, Haloferax volcanii. Eur J Biochem. 248(2):362–8.

Mevarech M, Frolow F, Gloss L. (2000) Halophilic enzymes: proteins with a grain of salt. Biophys Chem. 86(2-3):155–64.

Munawar N & Engel P (2013) Halophilic enzymes: characteristics, structural adaptation and potential applications for biocatalysis. Curr. Biotech. 2(4) 334–344

Nath A (2016) Insights into the sequence parameters for halophilic adaptation. Amino Acids. 48(3):751–762.

Overman L, Bujalowski W, Lohman T (1988) Equilibrium binding of Escherichia coli single-strand binding protein to single-stranded nucleic acids in the (SSB)65 binding mode. Cation and anion effects and polynucleotide specificity. Biochemistry 1988, 27(1) 456–471

Park C & Raines R (2001) Quantitative Analysis of the Effect of Salt Concentration on Enzymatic Catalysis. J Am Chem Soc. 123(46):11472–9.

Park C & Raines R (2003) Catalysis by Ribonuclease A Is Limited by the Rate of Substrate Association. Biochemistry 42(12):3509–3518

Patel S, Saraf M. (2015) Perspectives and Application of Halophilic Enzymes. In: Maheshwari D., Saraf M. (eds) Halophiles. Sustainable Development and Biodiversity, vol 6. Springer.

Pickl A & Schönheit P (2015) The oxidative pentose phosphate pathway in the haloarchaeon Haloferax volcanii involves a novel type of glucose-6-phosphate dehydrogenase - The archaeal Zwischenferment. FEBS Lett 589(10):1105–11.

Plantinga M, Korennykh A, Piccirilli J, Correll C (2008) Electrostatic Interactions Guide the Active Site Face of a Structure-Specific Ribonuclease to Its RNA Substrate. Biochemistry. 2008 Aug 26; 47(34):8912–8918.

Record Jr M, Lohman M, Haseth P (1976) Ion effects on ligand-nucleic acid interactions. J Mol Biol. 107(2):145–58.

Song B, Cho J, Raleigh D (2007) Ionic-strength-dependent effects in protein folding: analysis of rate equilibrium free-energy relationships and their interpretation. Biochemistry 46(49):14206–14.

Wright D, Banks D, Lohman J, Hilsenbeck J, Gloss L (2002) The effect of salts on the activity and stability of Escherichia coli and Haloferax volcanii dihydrofolate reductases. J Mol Biol. 323(2):327–44.

