## Supplementary material for "Unveiling the influence of salt concentration on the different stages of the catalytic cycle of a halophilic enzyme"

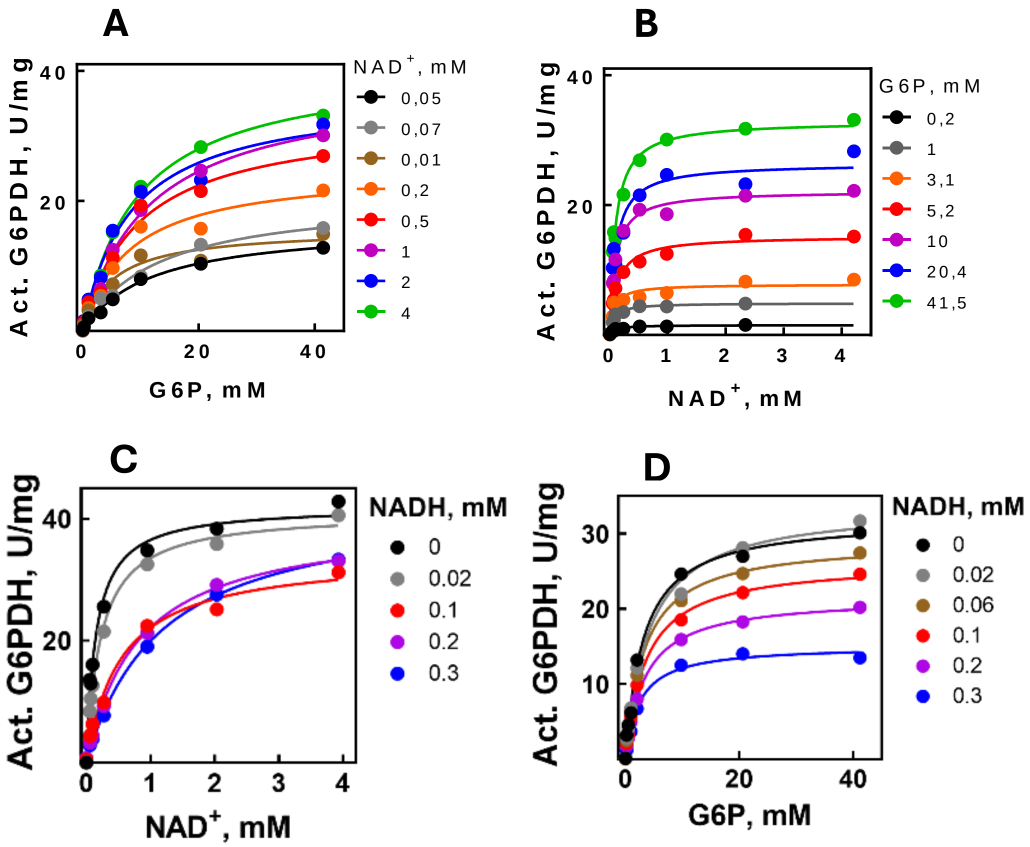


**Figure 1. Initial velocity experiments varying cosubstrate and product concentrations to determine the kinetic mechanism**.

A) saturation curves of G6P performed at different $NAD^+$ concentrations. B) saturation curves of $NAD^+$ performed at different G6P concentrations. C) Saturation curves for G6P obtained at different NADH concentrations. D) Saturation curves for $NAD^+$ obtained at different NADH concentrations. For each curve the Michaelis-Menten equation was fitted to each curve by non-linear regression.

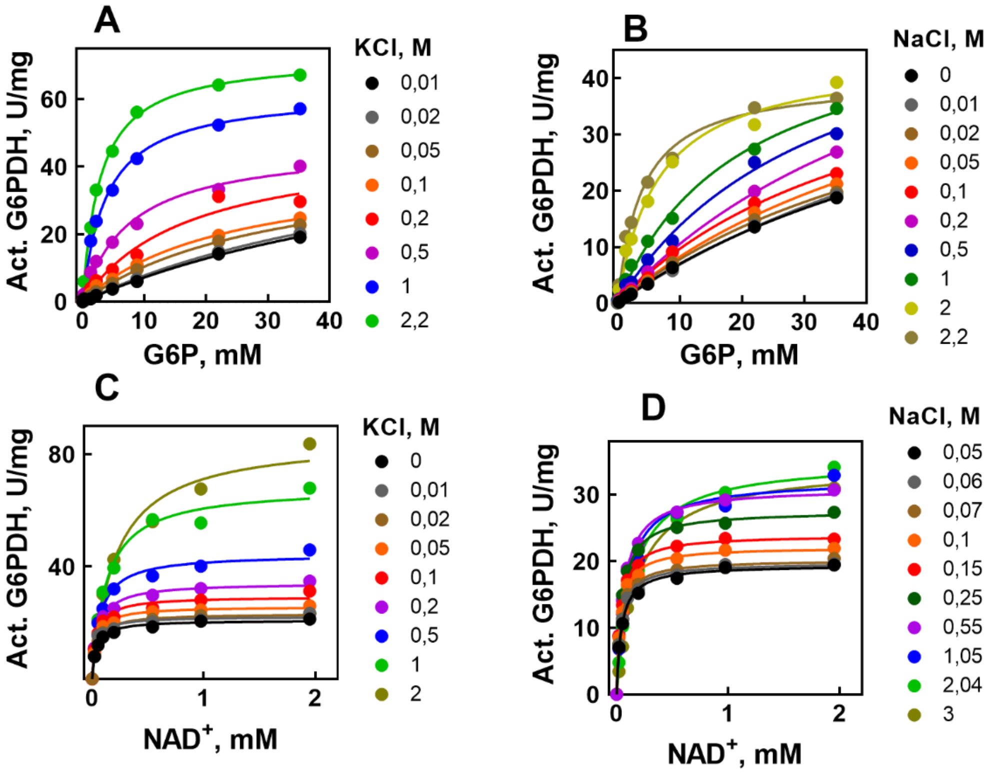


**Figure S2. Saturation curves for** G6P and $NAD^+$ at different **salt concentrations.**

Saturation curves for G6P were performed at different KCl (A) and NaCl (B) concentrations. curves for $NAD^+$ were performed at different KCl (F) and NaCl (G) concentrations. The Michaelis-Menten equation was fitted to each curve. All experiments were performed at 25 °C in Tris pH 8.5.

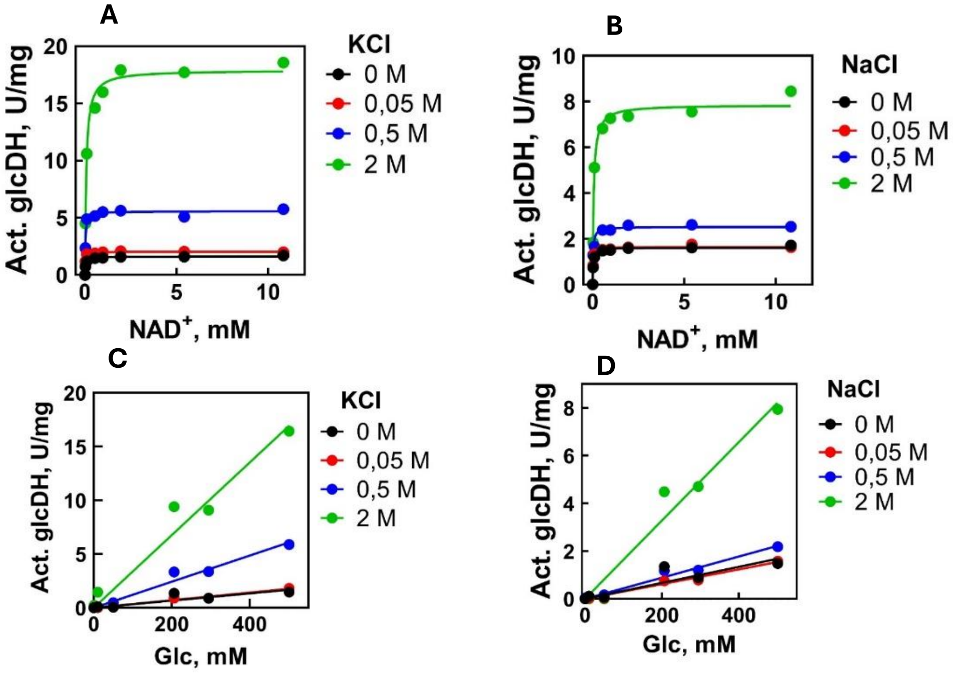


**Figure S2. Saturation** curves for $NAD^+$ and **glucose at different salt concentrations.**

Saturation curves for NAD+ were performed at different KCl (A) and NaCl (B) concentrations. The Michaelis-Menten equation was fitted to each curve. curves for glucose were performed at different KCl (F) and NaCl (G) concentrations. A straight line was fitted to each curves. All experiments were performed at 25 °C in Tris pH 8.5.
